# Environmental specificity and evolution in *Drosophila*-bacteria symbiosis

**DOI:** 10.1101/546838

**Authors:** Robin Guilhot, Antoine Rombaut, Anne Xuéreb, Kate Howell, Simon Fellous

## Abstract

Environmentally acquired microbial symbionts could contribute to host adaptation to local adaptation like vertically transmitted symbionts do. This scenario necessitates symbionts to have different effects in different environments. In *Drosophila melanogaster*, communities of extracellular bacterial symbionts vary largely among environments, which could be due to variable effects on phenotype. We investigated this idea with four bacterial strains isolated from the feces of a *D. melanogaster* lab strain, and tested their effects in two environments: the environment of origin (i.e. the laboratory medium) and a new one (i.e. fresh fruit with live yeast). All bacterial effects on larval and adult traits differed among environments, ranging from very beneficial to marginally deleterious. The joint analysis of larval development speed and adult size further suggests bacteria would affect developmental plasticity more than resource acquisition in males. The context-dependent effects of bacteria we observed, and its underlying mechanisms, sheds light on how environmentally acquired symbionts may contribute to host evolution.

## Introduction

Symbiosis contributes to host evolution through recruitment of adequate microorganisms (Margulis & Fester 1991; Jaenike et al. 2010; Fellous et al. 2011). As the environment varies among localities, different symbionts may be most beneficial in different conditions, possibly explaining microbiota variation among populations of the same animal species (e.g. Chandler et al. 2011; McKenzie et al. 2017). Microbial symbionts may therefore participate to local adaptation (Kawecki and Ebert 2004). A necessary condition to symbiont-mediated local adaptation is that microbial effects on host fitness change with environmental conditions (Schwab et al. 2016; Callens et al. 2016). The determining of host phenotype by interactions between symbiont identity and environment (i.e. Symbiont-by-Environment interactions) would thus largely be similar to so-called Genotype-by-Environment interactions that underlie genome-based local adaptation. Most studies exploring symbiont-mediated local adaptation have focused on vertically transmitted microorganisms (e.g. Moran et al. 2008). However, numerous animals form symbioses with bacteria that are in part acquired from the environment either by horizontal transmission between hosts or recruitment of free-living strains (Ebert 2013). Here, we explore how the effects of extracellular symbiotic bacteria on insect host traits change when hosts and bacteria are studied in an environment different from the one of origin.

*Drosophila* flies serve as important model organisms for host-microbiota studies (Douglas 2018). In *Drosophila melanogaster*, bacterial symbionts participate to a broad range of functions including resource acquisition, digestion, immunity and behavior (Broderick and Lemaitre 2012; Ankrah and Douglas 2018; Schretter et al. 2018). Several laboratory studies have established fly nutrition relies on interactions with gut bacteria (Shin et al. 2011; Storelli et al. 2011; Ridley et al. 2012; Wong et al. 2014; Huang et al. 2015; Leitão-Gonçalves et al. 2017; Téfit et al. 2017). In particular, bacterial genera frequently associated with laboratory flies, such as *Lactobacillus* and *Acetobacter*, can improve larval growth and development when laboratory food is poor in proteins (Shin et al. 2011; Storelli et al. 2011; Téfit et al. 2017). Even though some bacterial taxa are frequent in laboratory colonies, the composition of *Drosophila* bacterial gut communities largely varies among laboratories (Chandler et al. 2011; Staubach et al. 2013; Wong et al. 2013; Vacchini et al. 2017). Studies have shown that bacterial microbiota composition appears to be determined by the laboratory where the Drosophila flies were reared more than by their species (Chandler et al. 2011; Staubach et al. 2013), demonstrating these symbionts are largely acquired from the fly environment. Empirical studies have nonetheless shown pseudo-vertical transmission of bacteria from mothers to offspring also occurs in the laboratory (Bakula 1969; Ridley et al. 2012; Wong et al. 2015; Téfit et al. 2018). Microbiota composition differences between laboratory and field flies have led authors to argue that symbiotic phenomena as observed in the laboratory may not reflect those occurring in natural conditions (Chandler et al. 2011; Winans et al. 2017). The laboratory and natural environments of *D. melanogaster* flies indeed differ in several aspects. The most striking difference may have to do with the composition of the nutritive substrate upon which the adults feed, copulate, oviposit and within which larvae develop. Indeed, wild flies reproduce on and in fresh or decaying fruit flesh, usually colonized by yeast, whereas laboratory flies are reared on an artificial, jellified and homogeneous diet that contains long-chained carbohydrates (e.g. starch), agar, preservatives and dead yeast cells or yeast extract. To this date, very few studies have investigated *Drosophila*-bacteria interactions in conditions comparable to those of the field. It is therefore unknown whether fly-bacteria interactions that occur in the laboratory are maintained in natural substrate.

We experimentally studied the symbiosis between a laboratory strain of *D. melanogaster* and four of its bacterial symbionts in laboratory conditions and in grape berries where we mimicked natural egg and bacterial deposition from mothers. The four bacteria were isolated from the feces of adult flies and chosen for their ease of cultivation and recognition on standard microbiological medium. After inoculating bacteria-free eggs with these four bacterial isolates, we recorded phenotypic fly traits at the larval and adult stages. Our results show drastically different effects of symbionts on the hosts in laboratory medium and natural substrate. Some differences among environments can be explained by the environment-specific mechanisms of bacterial benevolence. The joint analysis of larval development time and adult size further suggests bacteria affect host developmental plasticity more than resource acquisition.

## Materials and Methods

### Drosophila strain

All insects were from the Oregon-R *Drosophila melanogaster* strain. This strain was founded in 1927 and has since been maintained in the laboratory. Our sub-strain was obtained from colleagues and reared on a laboratory medium comprising banana, sugar, dead yeast, agar and a preservative (Table S2.1). Before and during the experiment reported here, animals were maintained at 21 °C (stocks) or 23°C (experiment), with 70% humidity and a 14h photoperiod.

### Microbial isolates

We isolated a small number of symbiotic bacterial strain from the flies. Our aim was to use bacteria that were easy to culture and recognize morphologically but not to sample the whole community of bacteria associated with our flies stock. An important choice was to focus on aerobic bacteria that grow rapidly on standard agar plates at 25 °C, which excluded the anaerobes *Lactobacillus* that are among the best known symbionts of *D. melanogaster*.

In order to isolate bacteria present in fly feces, several groups of twenty *Drosophila melanogaster* flies were placed in sterile glass vials for 1 h. After fly removal, vials were washed with sterile PBS (Phosphate-Buffered Saline) solution, which was then plated on Lysogeny Broth (LB) agar medium (Table S2.2) and incubated at 24 °C. Four bacterial morphotypes were chosen based on visible and repeatable differences in size, color, general shape and transparency during repeated sub-culturing on fresh media (Figure S3). A single colony of each morphotype was amplified in liquid LB medium in aerobic conditions at 24 °C for 72 h, centrifuged and washed in PBS. Several sub-samples of equal concentration were stored at −80 °C in PBS with 15% glycerol and further used in this experiment (one per experimental block).

Molecular identification of each bacterium was carried out with Sanger sequencing. To this aim, a fresh colony of each bacterial type was picked with a sterile toothpick and dipped into sterile water, then boiled 10 min at 95 °C (Mastercycler, Eppendorf) and cooled in ice water. A sterile toothpick dipped into sterile water served as sterility control of the process. Fragments of the 16sRNA gene were amplified with bacterial primers Y2MOD (5-ACTYCTACGGRAGGCAGCAGTRGG-3’) and 16SB1 (5’-TACGGYTACCTTGTTACGACTT-3’) (Haynes et al. 2003; Carletto et al. 2008). PCRs were performed in a volume of 25 µl, containing each primer at 0.2 µM, 1x buffer (containing 2 mM MgCl_2_), each dNTP at 0.2 mM, and 1 U of *DreamTaq* Taq (Thermo Scientific). PCRs cycles had an initial denaturation step at 95 °C for 15 min, followed by ten cycles at 94 °C / 40 s - 65 °C / 45 s – 72 °C / 45 s); followed by 30 cycles at 94 °C / 40 s – 55 °C / 45 s – 72 °C / 45 s; and finished with an extension step of 10 min at 72 °C. Negative PCR controls were included. PCR products were visualized under UV light in an agarose gel before sequencing. Consensus sequences were created with CodonCode Aligner 4.2.7. Online SINA alignment service (https://www.arb-silva.de/aligner/) (Pruesse et al. 2012) and NCBI GenBank blastn service (https://blast.ncbi.nlm.nih.gov/Blast.cgi) were used to compare and assign the sequences. The four bacteria were identified as a *Staphylococcus* (likely *S. xylosus*), an *Enterococcus* (likely *E. faecalis*), an Enterobacteriaceae and an Actinobacteria (likely *Brevibacterium*). Further in this article, theses bacteria are referred to as *Staphylococcus*, *Enterococcus*, Enterobacteriaceae and Actinobacteria, respectively. All sequences were deposited in the NCBI database under the accession numbers MK461976 (*Staphylococcus*), MK461977 (*Enterococcus*), MK461978 (*Enterobacteriaceae*) and MK461979 (*Actinobacteria*).

A wild isolate of *Saccharomyces cerevisiae* yeast was used in experiments where larvae developed in fresh grape berries. The yeast was isolated from a wild Drosophilid in a vineyard in Southern France (*‘Le Domaine de l’Hortus’*, Hérault, France) (see Hoang et al. (2015) for a balanced discussion on *Drosophila-Saccharomyces* interactions). The isolate was grown in YPD medium, washed, split into several samples, stored at −80 °C in sterile PBS with 15% glycerol, that were further used in the experiment (one per block).

### Experimental design

We followed a full-factorial design resulting in twelve different treatments to assay: i. two types of fly environments - laboratory medium and grape berry (white, unknown cultivar) -, ii. six different symbiont treatments - the four bacterial strains described above, a mix of these four bacteria and controls without added bacteria. Each treatment had 13 to 15 replicates organized in 15 blocks launched over four days. Bacterial growth was also studied in fly-free grapes but is not described here.

Grape berries were surface-sterilized with 2% bleach solution before use. Because *D. melanogaster* females only oviposit in wounded fruit, we incised 5mm of berry skin (Figure S4) where we deposited twenty eggs free from culturable bacteria. These eggs were produced by the oviposition of flies on laboratory medium supplemented with the antibiotic streptomycin (1 mg / ml in 1 mM EDTA, Sigma-Aldrich ref. 85886). The efficacy of this method for removing culturable bacteria from egg surface was confirmed by the lack of bacterial growth after the deposition of such eggs onto LB agar plates (note however that detection of anaerobic bacteria such as *Lactobacillus* was not feasible in such conditions). Grape berries were inoculated with live yeast cells as it is a key component (Begg & Robertson 1948; Becher et al. 2012) and was necessary for fly survival in our system (Figure S1). For treatments with laboratory diet we deposited 20 eggs free from culturable bacteria on incisions at the surface of 4 ml of medium placed in 2 cm * 2 cm plastic cubes. Berries and laboratory media were all placed in 75 ml plastic vials closed by a foam plug.

Bacterial cells were inoculated to laboratory medium and grape berry immediately before egg deposition. Single bacterial strain treatments received 2.5 x 10^3^ live bacterial cells, and the mixed treatment 2.5 x 10^3^ cells of each bacterium (giving 10^4^ cells in total), suspended in 10 µl of sterile PBS. The number of inoculated bacterial cells, that is < 10^4^ Colony Forming Units (CFUs), was chosen based on the average number of bacteria previously reported in the guts of second-instar larvae (Bakula 1969; Storelli et al. 2011). In control treatments, sterile PBS was deposited instead of bacteria. On grape berries, 10^4^ live cells of the yeast *Saccharomyces cerevisiae* were inoculated. Note fruit substrate and live yeast presence are confounded factors in our experiment because we did not intend to study the effect of live yeast onto larval growth (Becher et al 2012) but to mimic field conditions where larvae develop in presence of live yeast. Although the laboratory medium also contains yeast, this preparation is inactivated (Table S2.1).

### Fly phenotyping

We recorded six different phenotypic traits in larvae and adults: larval size, larval mouthpart movement speed, number of larvae visible on medium surface, survival until adult emergence, time until adult emergence and a proxy of adult size. Larval traits were measured five days after egg deposition using a stereomicroscope. Larval mouthpart movement speed is the number of back-and-forth movements of the mouthpart that could be observed in 5 seconds. Newly formed pupae were transferred to empty sterile vials daily. We recorded male and female emergences daily. The size of adults, and their microbial content (see below), were estimated on a subset of those that emerged from the same vials. For each experimental replicate, we randomly selected a pupa before its emergence and when it emerged we pooled together all the flies of the same sex that emerged on the same day and from the same vial than the randomly selected pupae. These pools were homogenized in 200 µl of sterile PBS using a sterile pestle, divided in two sub-samples and stored at −80 °C with 15% sterile glycerol. One of the two sub-samples was used to numerate live bacteria and yeast cells in newly emerged adults, the other one to estimate adult size with the spectrophotometric method described in Fellous et al. (2018). We chose this method as it allowed the simultaneous analysis of adult size and microbial content. Briefly, we used log-transformed optical density at 202 nm of fly homogenate as a proxy of adult size. This was measured several months after the experiment when samples were thawed, crushed a second time using a Tissue Lyser II (Qiagen) for 30 s at 30 Hz with Ø3 mm glass balls, centrifuged for 30 s at 2000 G. Optical density of 15 µL of supernatant was then read on a Multiskan GO spectrometer (Thermo Scientific). This metrics correlates in both males and females with wet weight and wing length (all R^2^ > 0.8), two frequently used size proxies in *Drosophila* studies.

### Analysis of microbial development and evolution

The microbial content (i.e. bacteria and yeast) of newly emerged adults, as well as the microbial content of the laboratory media and of the grape berries after the removal of the last pupa were analyzed. In this manuscript we only report on the presence or absence of inoculated bacteria in the larval environment. We will describe the transmission of inoculated bacteria and yeast from larvae to adults (i.e. through metamorphosis) in a separate manuscript.

In order to better understand fly symbiosis with the Enterobacteriaceae and the Actinobacteria we analyzed their metabolic abilities profiles with Eco Microplates (Biolog) that contain 31 different carbon substrates (Text S5). In one case we recorded the presence of the Actinobacteria in a grape berry at the end of the experiment. The bacterium was isolated and its metabolic abilities were compared to that of its ancestor deposited at the beginning of the experiment. Because we suspected the grape-retrieved Actinobacteria had evolved the ability to better develop in fruit flesh we compared its growth in grape flesh to that of the ancestor (Text S6). The two bacteria were deposited in slices of surface-sterilized berries with two initial concentrations 10^4^ cells and 10^6^ cells (eight replicates of each). Grape disks were sampled after 24 h and 72 h and bacteria numerated on LB agar plates.

### Data analysis

To study the response of fly phenotypes to variation of larval substrate and bacterial symbiont, linear mixed models (LMM) with Restricted Maximum Estimate Likelihood (REML) were used. ‘Block identity’ was defined as random factor, while we defined as fixed factors the ‘larval environment’ (i.e. laboratory medium or fruit), ‘bacterial treatment’, ‘fly sex’ (for the analyses of age at emergence and adult size only), and their full-factorial interactions. Homoscedasticity and residuals normality complied visually with model assumptions. Post-hoc Student’s tests were used to decipher significant differences among factor levels.

Bacteria and fungi different from those we inoculated were observed in 17% of the vials, which were further excluded for all analyzes presented in this article. Results of fly traits analyses were identical in the full and the curated dataset. Both datasets are available online.

The number of Actinobacteria cells (counted by colony forming units CFUs) in grape disks inoculated with the ancestral strain and the isolate retrieved from a replicate of the experiment was analyzed with a linear model. Number of cells was log(x+1) transformed to comply with model assumptions. The full-factorial model contained the factors ‘bacterial strain identity (ancestor or derived)’, ‘time after inoculation (24 h or 72 h)’ and ‘initial cell concentration (10^4^ or 10^6^ cells)’. Post-hoc tests were carried out by comparing with linear models cell numbers of the ancestor and derived isolate in each of the four combinations of initial density and time after inoculation. The metabolic abilities of the ancestral and derived Actinobacteria - as assayed with Eco Microplates (Text S5) in three independent observations per substrate, bacterial strain and assay duration - were further compared with Mann-Whitney tests for each individual carbon source.

Analyzes were performed with JMP (SAS, 14.1) and R (version 3.5.2).

Dataset is available in the open data repository Zenodo (DOI: 10.5281/zenodo.2554194).

## Results

### Larval traits

*Larval size* after five days was influenced by an interaction between the environment and the bacterial treatment (Table 1, Figure 1A). In grapes, the Actinobacteria decreased larval size relative to bacteria-free controls but had no particular effect in laboratory media. In laboratory media, the Enterobacteriaceae produced large larvae both alone and when mixed with the other bacterial strains, which did not happen when grown on a grape substrate.

**Table 1:**
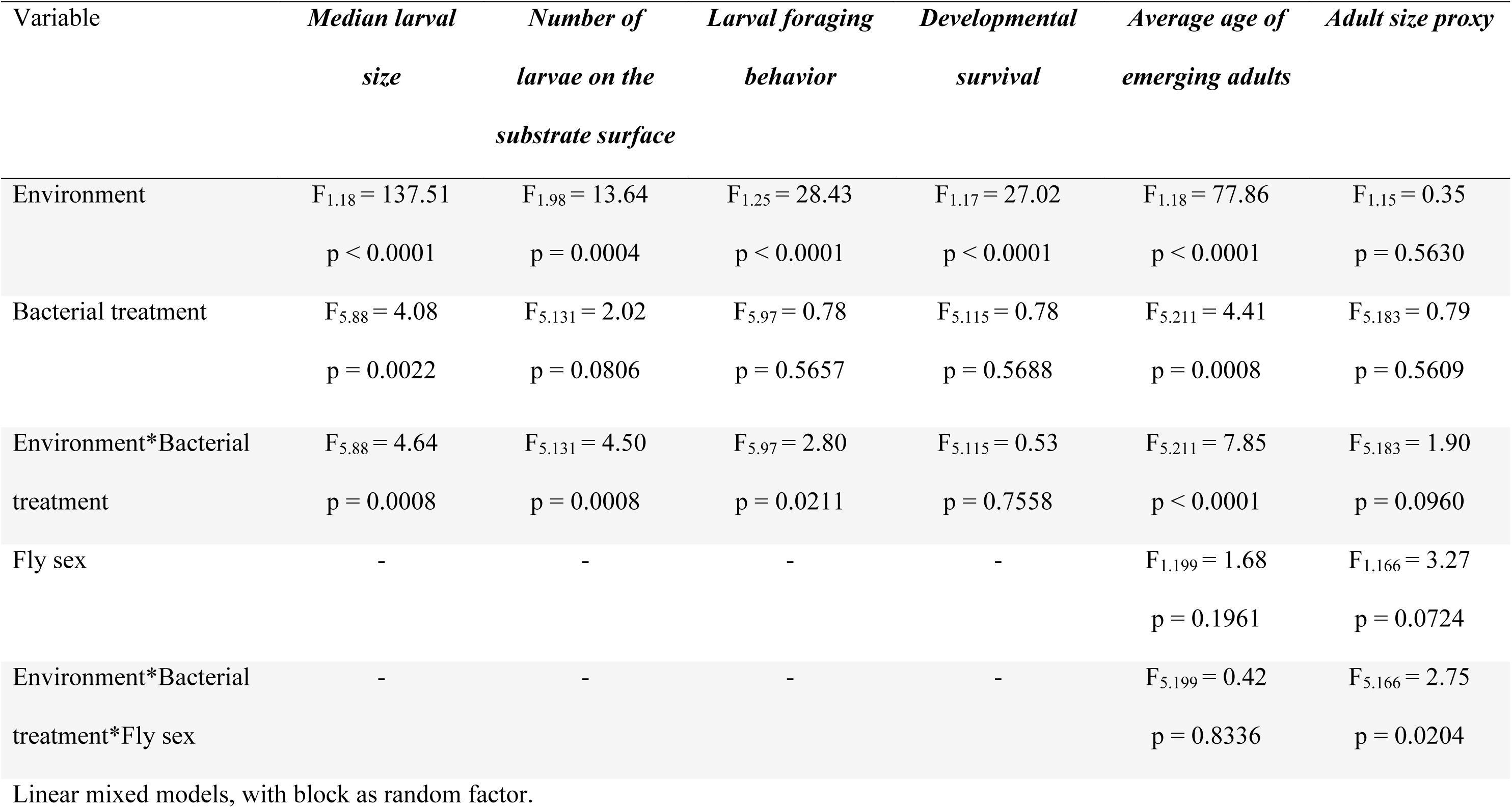
analysis of larval and adult phenotypes in response to bacterial treatment and larval environment. Linear mixed models (REML).

| Variable | *Median larval*<br>*size* | *Number of*<br>*larvae on the*<br>*substrate surface* | *Larval foraging*<br>*behavior* | *Developmental*<br>*survival* | *Average age of*<br>*emerging adults* | *Adult size proxy* |
| --- | --- | --- | --- | --- | --- | --- |
| Environment | $F_{1,18} = 137.51$<br>$p < 0.0001$ | $F_{1,98} = 13.64$<br>$p = 0.0004$ | $F_{1,25} = 28.43$<br>$p < 0.0001$ | $F_{1,17} = 27.02$<br>$p < 0.0001$ | $F_{1,18} = 77.86$<br>$p < 0.0001$ | $F_{1,15} = 0.35$<br>$p = 0.5630$ |
| Bacterial treatment | $F_{5,88} = 4.08$<br>$p = 0.0022$ | $F_{5,131} = 2.02$<br>$p = 0.0806$ | $F_{5,97} = 0.78$<br>$p = 0.5657$ | $F_{5,115} = 0.78$<br>$p = 0.5688$ | $F_{5,211} = 4.41$<br>$p = 0.0008$ | $F_{5,183} = 0.79$<br>$p = 0.5609$ |
| Environment*Bacterial<br>treatment | $F_{5,88} = 4.64$<br>$p = 0.0008$ | $F_{5,131} = 4.50$<br>$p = 0.0008$ | $F_{5,97} = 2.80$<br>$p = 0.0211$ | $F_{5,115} = 0.53$<br>$p = 0.7558$ | $F_{5,211} = 7.85$<br>$p < 0.0001$ | $F_{5,183} = 1.90$<br>$p = 0.0960$ |
| Fly sex | - | - | - | - | $F_{1,199} = 1.68$<br>$p = 0.1961$ | $F_{1,166} = 3.27$<br>$p = 0.0724$ |
| Environment*Bacterial<br>treatment*Fly sex | - | - | - | - | $F_{5,199} = 0.42$<br>$p = 0.8336$ | $F_{5,166} = 2.75$<br>$p = 0.0204$ |

Linear mixed models, with block as random factor.

**Figure 1:**
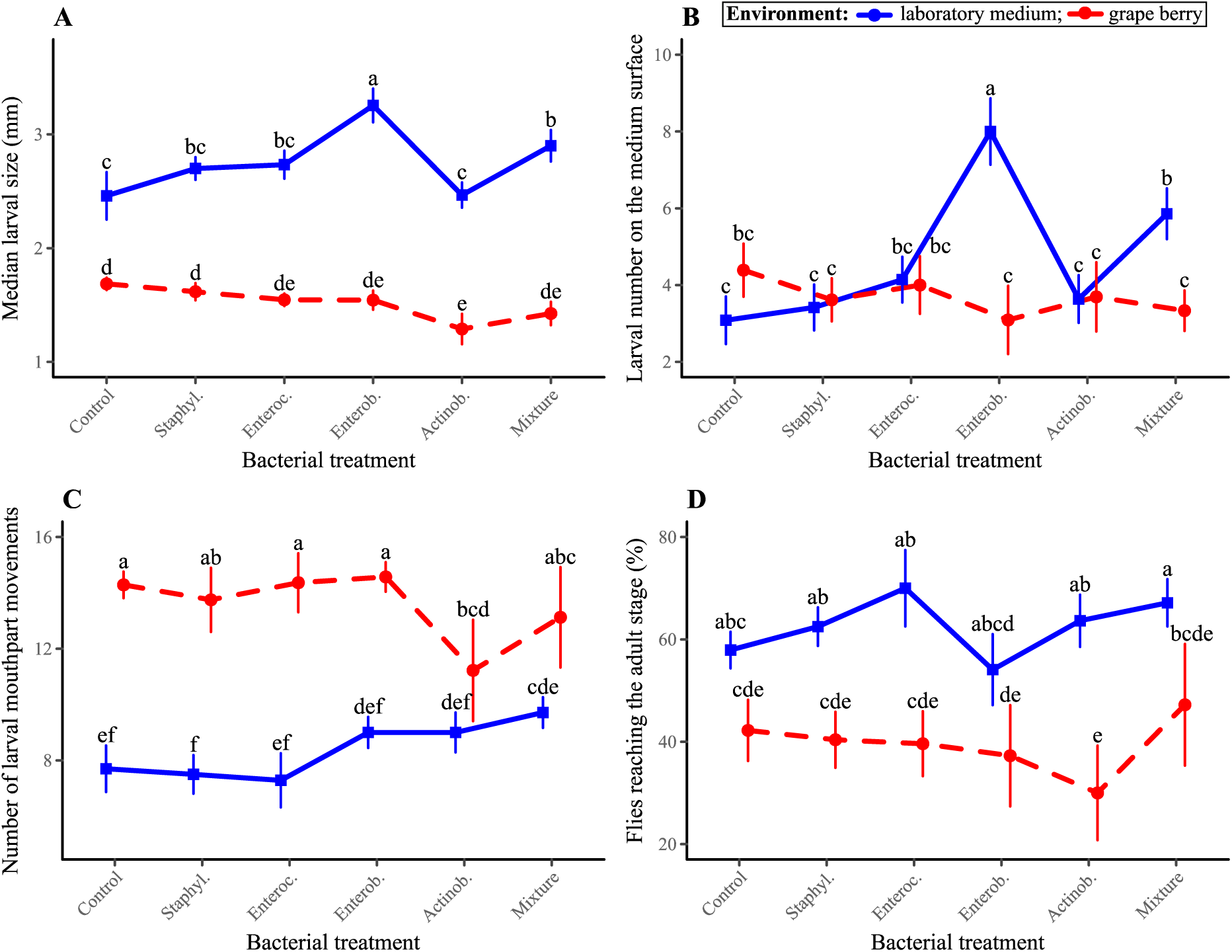
larval phenotypes in response to bacterial treatment and larval environment. (A) Median larval size; (B) Number of larvae on the medium surface; (C) Number of larval mouthparts movements; (D) Developmental survival. Symbols indicate means; error bars indicate standard errors around the mean. Means not connected by the same letters are significantly different.

*The number of larvae visible on medium surface* was influenced by an interaction between the environment and the bacterial treatment (Table 1, Figure 1B). Presence of the Enterobacteriaceae in laboratory media led to greater numbers of visible larvae compared to all other treatments.

*Mouthparts movement pace* was influenced by an interaction between the environment and the bacterial treatment (Table 1, Figure 1C). Movements were generally faster in grapes than in laboratory media. However, the Actinobacteria slowed down the movements of mouthparts in grapes to a level comparable to the one of larvae reared on laboratory media.

*The proportion of eggs surviving until the adult stage* was only affected by the environment, with a lower survival in grapes than in laboratory media (Table 1, Figure 1D).

### Metamorphosis and adult traits

*Age at adult emergence* was not different among sexes but influenced by an interaction between the environment and the bacterial treatment (Table 1, Figure 2A). In laboratory media, flies reared with the Enterobacteriaceae, alone or in mixture, emerged nearly two days sooner than other flies in the same environment and almost four days earlier than those in grapes (Figure 2A). No differences were observed among bacterial treatments in grapes (Figure 2A).

**Figure 2:**
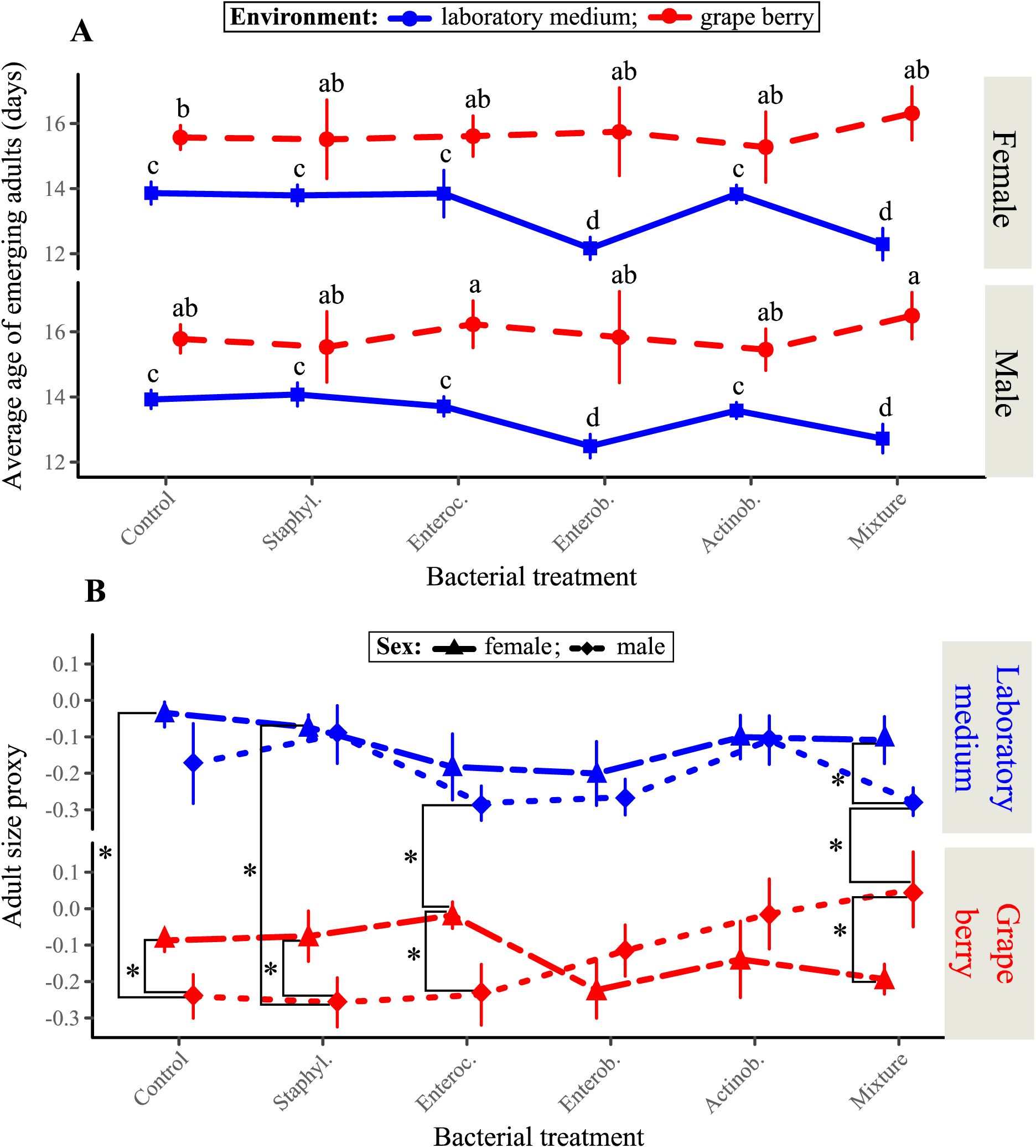
adult *Drosophila* phenotypes in response to bacterial treatment and larval environment. (A) Average age of emerging adult females and males; (B) Adult size proxy. Symbols indicate means; error bars indicate standard errors around the mean. Means not connected by the same letters (Figure A) or * (Figure B) are significantly different.

*Adult size* was influenced by a triple interaction between sex, the environment and the bacterial treatment (Table 1, Figure 2B). Several bacterial treatments had sex-specific effects that differed among environments. For example, the mixture of all four bacteria produced larger males than females in grapes but smaller males than females in laboratory media. Similarly, the *Staphylococcus* produced larger females in grapes and in laboratory media than males in grapes.

### Joint effect of bacteria on age at emergence and adult size

It is well established in numerous animals that, all else being equal, the speed of larval development (i.e. 1/ age at maturity) trade-offs with adult size (Teder et al. 2014). This gave us the opportunity to disentangle symbiont effects on resource acquisition (i.e. performance that can bolster one trait with no cost to the other) from developmental plasticity along the trade-off. To this end, we related developmental speed and adult size using the mean trait value of each bacterial treatment (Figure 3). It was mandatory to remove the overall effects of each environment on host phenotypes, otherwise, if one environment was generally more favorable than the other it would have created a positive relationship between larval development speed and adult size that could have concealed the influence of the bacteria on the plastic relationship between these traits. We therefore divided the mean trait values of each treatment (5 bacterial treatments * 2 environments = 10 treatments) by that of the bacteria-free controls in the same environment (grape or laboratory medium). Males and females were analyzed separately owing to the three-way interaction between sex, environment and bacteria observed for adult size.

**Figure 3:**
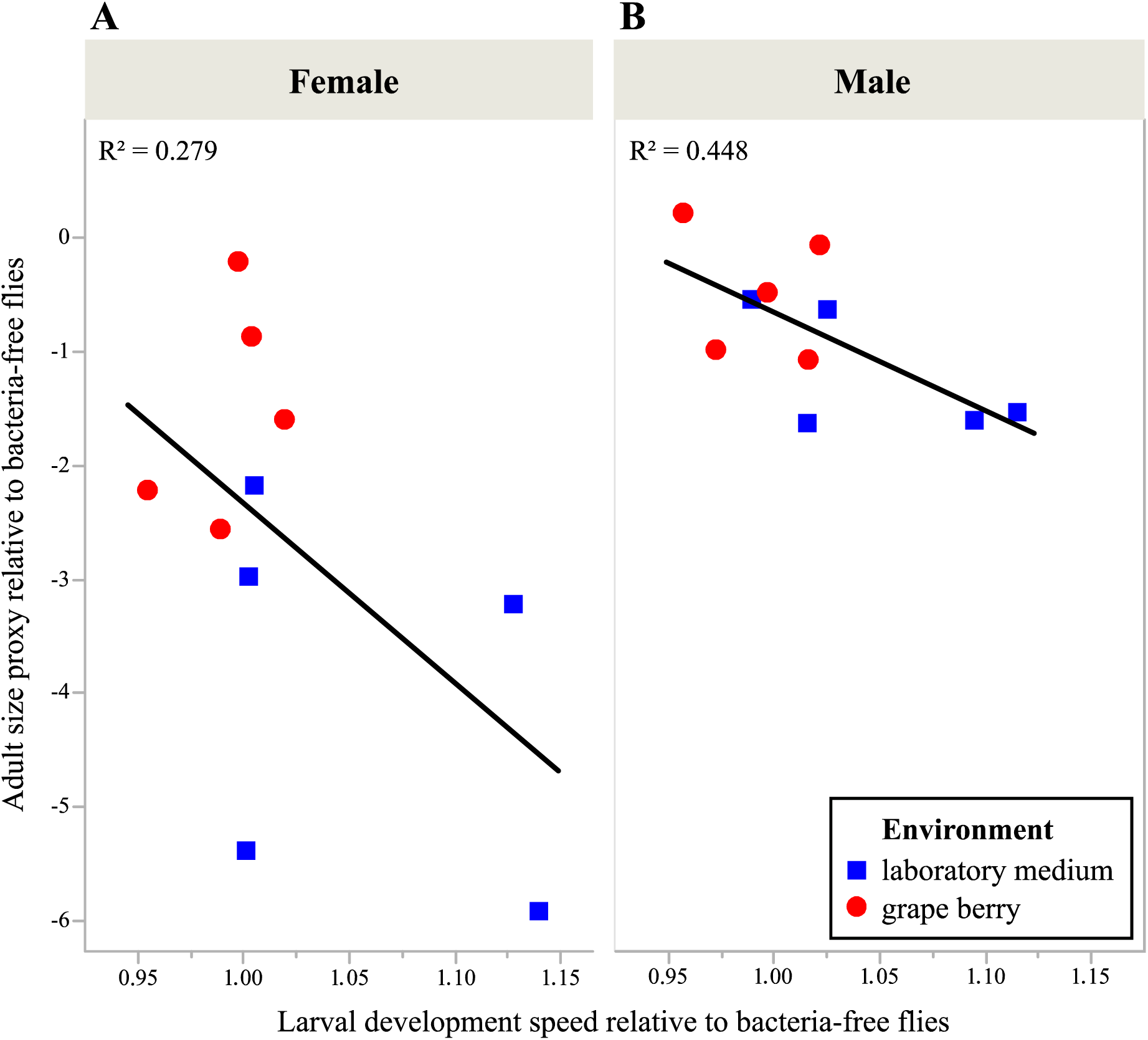
relationship between average effects of bacteria on larval developmental speed and adult size, in females (A) and females (B). Symbols indicate the phenotype means of each treatment (i.e. combination of bacterium and larval environment). All values are expressed relative to average observations of bacteria-free treatments.

The relationship between effects of bacteria on duration of larval stage and adult size was marginally significant and negative for males (Linear model F_1.8_ = 6.48, p = 0.0344) but not for females (F_1.8_ = 3.09, p = 0.1169) (Figure 3). Overall, in grapes bacteria produced slow-developing but large males, while they developed faster and were smaller in laboratory media (Figure 3B).

### Bacterial cells in the environment and the flies

The Enterobacteriaceae isolate was the only bacterium to be consistently retrieved from the environment in which larvae had developed, however only in laboratory media (Figure 4).

**Figure 4.**
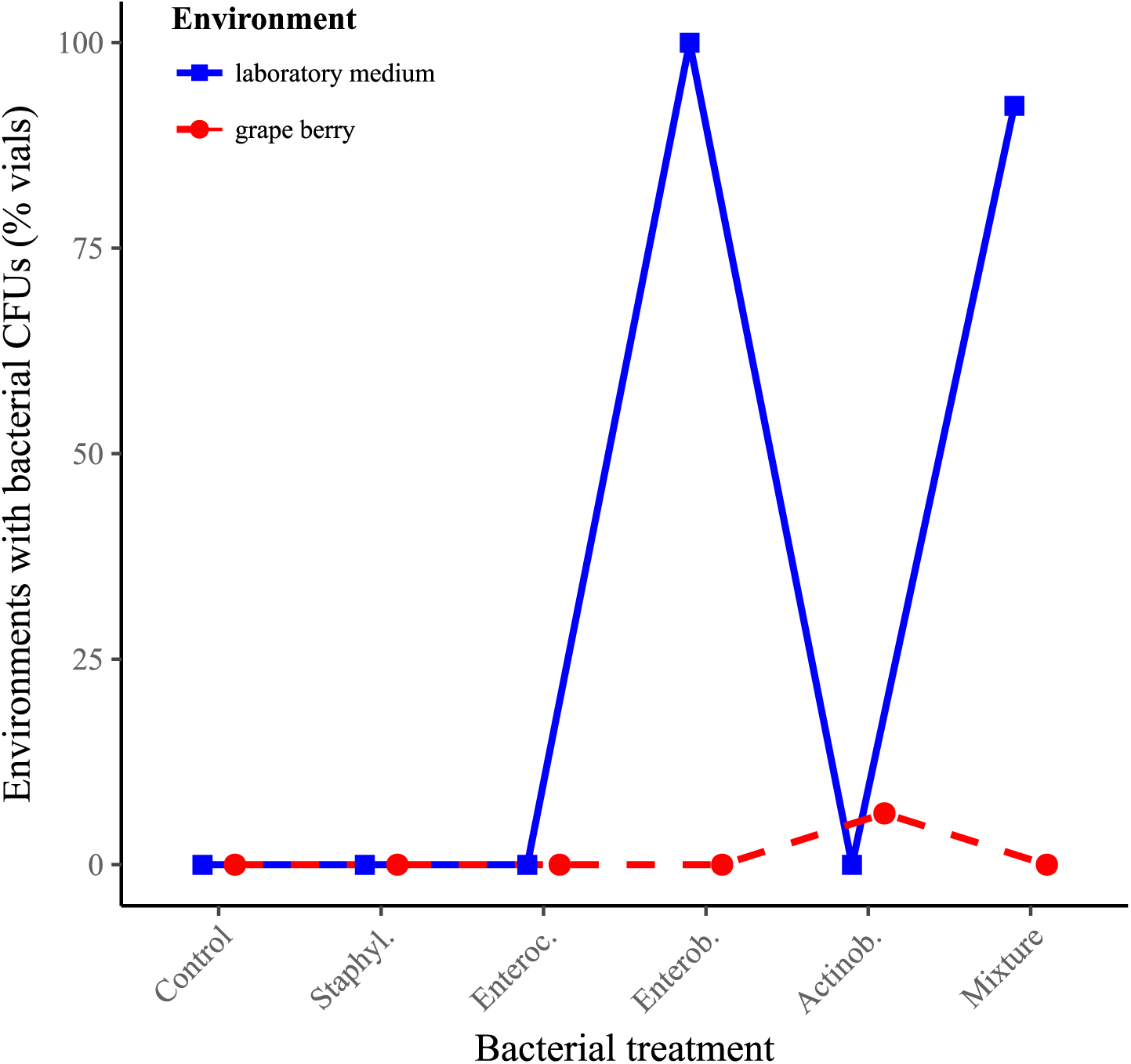
Proportions of environments containing bacterial cells of the strain inoculated as observed after the formation of the last pupa. Symbols indicate percentages per treatment.

In one instance, the Actinobacteria was found in a grape berry from which no live adult fly emerged. This bacterium was isolated and further studied in order to investigate its possible evolution (Text S6). Comparison between the Actinobacteria ancestor and the isolate from fruit revealed a marginally non-significant interaction between bacterium identity, sampling time and initial cell density (F_1.56_ = 3.25, p = 0.077) (Figure S6). Comparison between the two bacteria at all time points and initial cell densities (i.e. 4 combinations) revealed a single significant difference between the ancestor and the derived strain after 24 h with an initial density of 10^4^ cells per grape disk (t = −3.33, p = 0.005). The metabolic differentiation of the two bacteria was investigated on thirty-one carbon sources. After 48 h, growth of the derived Actinobacteria was significantly greater than that of the ancestor in eleven substrates, and lower in four substrates (Figure S6A). After 120 h, growth of the derived Actinobacteria was significantly greater than that of the ancestor in three substrates, and lower in sixteen substrates (Figure S6B).

## Discussion

We studied the symbiotic interactions between a laboratory strain of *Drosophila melanogaster* and four bacterial strains isolated from adult feces. No single effect of the bacteria on host phenotype observed in laboratory medium (i.e. the environment of origin) maintained in fresh fruit (i.e. the environment close to natural conditions). Some of these observations can be explained by the ecology of laboratory-associated symbionts in artificial medium. Further analyses suggest combination of environment and bacteria affected host developmental plasticity along a trade-off between larval growth speed and adult size.

### Different symbiont effects in different environments

The observation that all bacterial effects on host phenotype were different in laboratory medium and grape berry prompts the question of the reason behind this discrepancy. Focusing of the Enterobacteriaceae may shed light onto the ecologies of the symbiotic bacteria we isolated, and why they differed among environments.

In laboratory medium, inoculation of the Enterobacteriaceae induced greater larval size and accelerated larval development (Figures 1A and 2A). Besides, adults produced by larvae associated with the Enterobacteriaceae in laboratory medium were not significantly smaller than in other treatments. The bacterium hence accelerated larval growth. In its presence larvae remained at the surface of the medium where they could be observed in greater numbers than with all other treatments (Figure 1B), even though there were no mortality differences among them (Figure 1D). The Enterobacteriaceae was also the only bacterium to be retrieved from the medium after fly pupation (Figure 4). These last two elements suggest the bacterium serves as food: it would grow on medium surface and be consumed by grazing larvae. This idea is further supported by the visual observation that, in absence of larvae, media inoculated with the Enterobacteriaceae harbored white microbial growth on their surface (Figure S7). Along these lines, the wide metabolic spectrum of this bacterium (Figure S5.1) is congruent with a microorganism being a generalist that would extract resources from the medium, possibly transform nutrients (Ankrah and Douglas 2018; Sannino et al. 2018), and eventually concentrate them on medium surface. This phenomenon would be in many ways similar to that described by Yamada et al. (2015) where the yeast *Issatchenkia orientalis* extracts amino acids from agar-based laboratory medium and concentrates them on medium surface where adult flies harvest them. The physical nature of laboratory medium is very different from that of real fruit. In particular, the agar of laboratory medium permits the diffusion of simple nutrients and their absorption by bacteria and yeast. However, in fresh fruit nutrients are not free to diffuse but enclosed in cells, it is therefore understandable that the Enterobacteriaceae did not accelerate larval growth in fresh fruit. Larvae feeding on surface growing microorganisms may therefore be more common in the laboratory than in the field, where larvae are rare on the surface of fruits.

Bacteria-induced nutritional effects on *Drosophila* larvae and adults are frequently attributed to gut bacteria (Shin et al. 2011; Storelli et al. 2011; Broderick and Lemaitre 2012; Leitão-Gonçalves et al. 2017). It is well established that lactic and acetic acid bacteria, two taxa that were not investigated in our experiment, can promote larval growth upon nutrient scarcity (Shin et al. 2011; Storelli et al. 2011, Téfit et al. 2017). However, it is also well established that bacteria can affect *Drosophila* phenotype through signaling (Storelli 2011) as well as nutrient provisioning (Brownlie et al. 2009; Bing et al. 2018; Sannino et al. 2018). In most cases, these effects which were described from laboratory flies and in laboratory medium, are condition specific (Douglas 2018). Indeed, bacteria are often only beneficial when laboratory food has a low concentration in dead yeast (i.e. amino acids) (Shin et al. 2011; Storelli et al. 2011). Our results extend these previous observations as the *Staphylococcus*, the *Enterococcus* and the Actinobacteria we isolated and assayed here not only lost their beneficial effects when tested out of laboratory medium, but also acquired new effects. For example, in grape larval size was reduced by the Actinobacteria relative to bacteria-free controls (Figure 1A) showing symbionts can become costly when associated with host in a new environment.

Whether flies' symbiotic bacteria reside durably in fly guts or are constantly excreted in the environment and re-absorbed during feeding is still debated (Ma & Leulier 2018; Pais et al. 2018). It is not possible to further this debate with our data. However, we note that only one of our four bacterial isolates, the Enterobacteriaceae, was consistently retrieved from the larval artificial medium (Figure 4). By contrast, all isolates were found in adults produced by pupae that were separated from the larval environment before emergence (data presented in another forthcoming manuscript). These observations are congruent with the hypothesis that the *Staphylococcus*, the *Enterococcus* and the Actinobacteria we isolated are gut residents rather than grow in the medium. In the only case where we retrieved the Actinobacteria from fruit flesh, it is striking that the ability of this bacterium to maintain in fruit seemed to have evolved in the course of our experiment (Figure 4). This association between bacterium evolution and effects of host phenotype echoes the results of Martino et al. (2018) who showed *Lactobacillus* adaptation to food medium leads to greater benevolence. However, in our case adaptation of the Actinobacteria to fruit environment associated with greater cost to the host.

### Host developmental plasticity

It is well established that holometabolous insects, such as fruit-flies, must trade-off duration of larval development (i.e. age at maturity) with adult size (Teder et al. 2014, Nunney 1996). Figure 3 displays the effects of bacterial and substrate treatments on larval and adult traits relative to treatments without added bacteria. Host trait values for each bacterial treatment were divided by values measured in controls reared on the same substrate but without addition of bacteria. The relationship between speed of larval development and adult male size was marginally significant and negative. Data therefore suggests that bacterial treatments that slowed down development led to the production of larger adult males. Because data-points from fruit and artificial medium segregated in different parts of phenotypic space the results may be partly driven by the environment-specific effects of bacteria on hosts. The negative relationship came as a surprise as we expected nutritional symbionts to affect developmental speed and adult size either in an independent or similar fashion, which would have led to an absence or a positive relationship between these two traits, respectively. Correlated, positive effects of a nutritional symbiont on larval and adult traits were for example shown in yeast-*Drosophila* mutualism (Anagnostou et al. 2010; Bing et al. 2018). For example, the species of yeast *Metschnikowia pulcherrima* produce small adults that are also slow to develop (Anagnostou et al. 2010). Our results suggest that bacterial symbionts, such as the ones we studied here, could alter developmental plasticity in response to the ecological context. This hypothesis is congruent with the known effect *Lactobacillus plantarum* bacteria on host development mediated by hormonal changes (Storelli et al. 2011). Whether microbial symbionts influence hosts through variation of general vigor (Fry 1993) or developmental plasticity (two non-excluding possibilities) may change the evolutionary fate of the host-symbiont relationship. Indeed, symbionts that plastically alter phenotypes may be more dispensable that those providing functions host genomes are not capable of (Fellous and Salvaudon 2009). It could further be argued the fitness effect of altering developmental plasticity may depend on environmental context more than general improvement of resource acquisition (Chevin et al 2010). As such symbiont mediated effects on host plasticity is in line with the idea that many symbionts have context-dependent effect on the fitness of their host (e.g. De Vries et al. 2004; Duncan et al. 2010; Daskin & Alford 2012; Bresson et al. 2013; Callens et al. 2016; Cass et al. 2016). We are now pursuing further investigation to determine if, and when, bacterial and yeast symbionts affect host developmental plasticity rather than general performance in *Drosophila* flies.

### Symbiont-mediated evolution

A consequence of *Drosophila* bacterial symbionts having different effects in different environments is the possibility they participate to the fine-tuning of host phenotype to local conditions (Margulis & Fester 1991; Moran 2007; Sudakaran et al. 2017). The phenomenon is now well established in vertically transmitted symbionts of insects that protect their hosts from parasites. For example, populations of aphids exposed to parasitoids harbor protective *Hamiltonella* symbionts at greater frequency than parasitoid-free populations (Oliver et al. 2005). Similarly, in the fly *Drosophila neotestacea*, the spread of the bacterium *Spiroplasma* allowed hosts to evolve greater resistance to parasitic nematodes (Jaenike et al. 2010). Vertically-transmitted bacterial symbionts of *Paramecium* ciliates can also improve host resistance to stressful conditions (Hori & Fujishima 2003). Whether bacteria act as parasites or mutualists then depends on the genetic ability of the host to deal with stress in absence of the symbiont (Duncan et al. 2010). However, the evolutionary role of symbionts that may be acquired from the environment is less clear, in part because the mechanisms favoring the association of hosts with locally beneficial symbionts are not as straightforward as for vertical transmission (Ebert 2013). Nonetheless, several lines of evidence suggest environmentally acquired microbial symbionts may participate to local adaptation in *Drosophila*-microbe symbiosis. First, symbionts can be transmitted across metamorphosis (i.e. transstadial transmission from the larval to the adult stage) and pseudo-vertically during oviposition (i.e. from mothers to offspring) (Bakula 1969; Starmer et al. 1988; Spencer 1992; Ridley et al. 2012; Wong et al. 2015; Téfit et al. 2018). Second, host immune system participates to the destruction of harmful gut bacteria and the retention of beneficial ones (Lee et al. 2017; Lee et al. 2018). Third, *Drosophila* larvae actively search and associate with beneficial yeast species ensuring they engage in symbiosis with locally adequate nutritional symbionts (Fogleman et al. 1981; Fogleman et al. 1982). In addition to preferential association with beneficial microbes, *Drosophila* adaptation to local conditions thanks to microorganisms further necessitates symbionts have different effects in different environments. Our results show bacteria isolated from a fly population can either be beneficial, neutral or costly depending on the substrate larvae were reared in (Figures 1 and 2). Bacterial symbionts may therefore participate to host adaptation in *Drosophila*-bacteria symbioses through variations in symbiont community composition.

Host adaptation based on symbionts differs from genome-based evolution in that microbes can provide a greater amount of evolutionary novelty than mutations of nuclear genes do (Jaenike et al. 2012; Moran 2007). This arises from several factors. A single metazoan individual can associate with billions of microbial cells that each has a genome with potentially beneficial mutations. It results that populations of microbial symbionts can adapt faster to local conditions than nuclear genes. *Caenorhabditis elegans* nematodes host a diversity of bacteria, some of which may be detrimental. A recent study demonstrated how rapid evolution in the competition between two bacterial species, one of which being pathogenic to worms, lead to host protection against the most virulent bacterium (King et al. 2016). Rapid symbiont evolution can also be beneficial to hosts in the case of nutritional symbioses as demonstrated in the relationship between *Drosophila melanogaster* and the bacterium *Lactobacillus plantarum*. It was recently shown that bacterium adaptation to nutritional substrate during 2000 generations (i.e. 313 days) in absence of hosts not only improves bacterial performance but also that of *Drosophila* larvae associated to the evolved bacterium (Martino et al. 2018). Our data suggest the pace of microbial evolution to environmental conditions may be even faster. Indeed, at the end of our experiment, we retrieved live Actinobacteria cells from one fruit. Preliminary experiments had shown this strain we had isolated from fly feces was not able to grow in grape flesh (Figure S6). We therefore hypothesized the Actinobacteria isolate had evolved a better ability to maintain in fruit flesh than the ancestor we had inoculated. Comparison between this derived strain and the ancestor indeed suggests the bacterium evolved better persistence in the environment in the time course of our experiment (Fig S6.1). However, conclusion based on this observation must not be over-stretched as our experimental setup was not initially designed to test for bacterial adaptation, we only observed this phenomenon once and the comparison between the ancestral and the derived strain is contingent on minute experimental details. On the other hand, the metabolic abilities of the derived isolate had evolved relative to the ancestor in the majority of the 31 carbon substrates they were tested on (Figure S5.2), suggesting rapid bacterial evolution did occur. The derived strain was collected in one of the replicates where larvae were smallest and with slowest mouthpart movements, where live yeast concentration was lowest, and from which no adult emerged, showing that bacterial adaptation to environmental conditions may be detrimental to insect hosts.

## Conclusion

In this study, we found that associations between laboratory *Drosophila* flies and their microbial symbionts result in different effects on host phenotype when the symbiosis is investigated under conditions close to nature. The context-dependence of bacterial effects, and the underlying mechanisms we unveiled (i.e. bacterial ecology, bacterial effects on host plasticity and rapid bacterial evolution), shed light on the role of microorganisms in the evolution of their hosts. Understanding the ecology and evolution of symbiosis in the wild will necessitate working with wild strains of animals and symbionts under ecologically realistic conditions, which is attainable in the *Drosophila* system.

## Acknowledgements

We warmly thank L. Benoit, M.P. Chapuis, D. Duneau, R. Gallet, P. Gautier, F. Leulier and N. Rode for methodological help and useful comments on an earlier version of this work.

## Supplementary Material 1. Live yeast as a prerequisite to *D. melanogaster* larvae survival on pristine grape berry

**Figure S1:**
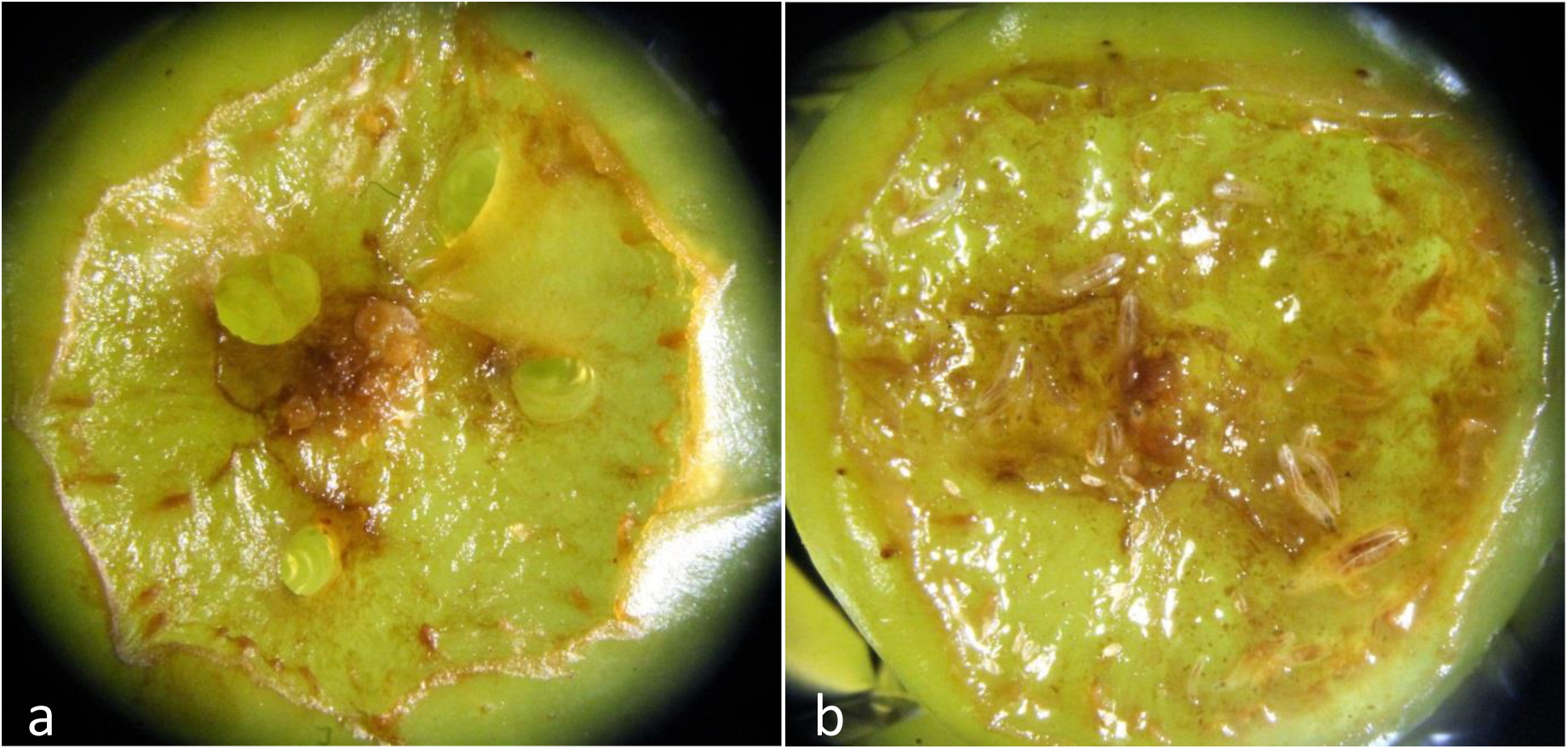
Live yeast is necessary for the survival of *D. melanogaster* larvae on pristine grape berry. Prior to the experiment, we investigated survival of *D. melanogaster* larvae on fresh grape berries. Twenty bacteria-free *D. melanogaster* eggs were deposited next to an artificial wound with or without the bacterial isolates and *Saccharomyces cerevisiae*. In absence of yeast, larvae died quickly after hatching, with or without bacteria (Figure S1a). When live yeast was added to the system, numerous larvae developed up to the 3^thrd^ instar (Figure S1b), when we stopped monitoring.

## Supplementary Material 2. Laboratory recipes

**Table S2.1:** Laboratory medium recipe.

| Component | *Amount for 1.5L* |
| --- | --- |
| Reverse osmosis water | 1200 ml |
| Banana | 280 g |
| Sugar | 74 g |
| Dead yeast | 74 g |
| Alcohol | 30 ml |
| Agar | 12 g |
| Nipagin | 6 g |

**Table S2.2:**
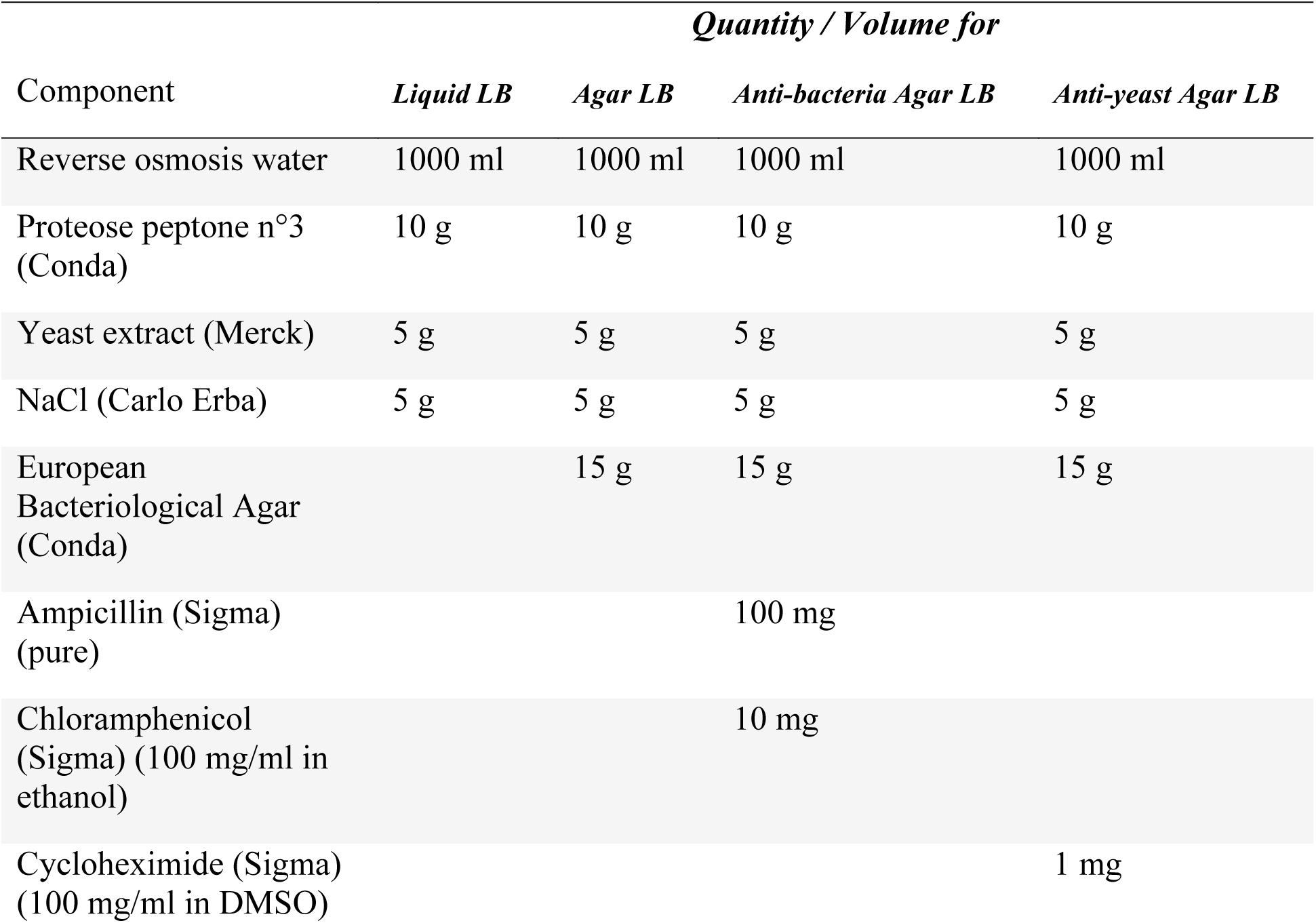
Lysogeny broth (LB) recipes.

| Component | *Quantity / Volume for* |  |  |  |
| --- | --- | --- | --- | --- |
|  | *Liquid LB* | *Agar LB* | *Anti-bacteria Agar LB* | *Anti-yeast Agar LB* |
| Reverse osmosis water | 1000 ml | 1000 ml | 1000 ml | 1000 ml |
| Proteose peptone n°3 (Conda) | 10 g | 10 g | 10 g | 10 g |
| Yeast extract (Merck) | 5 g | 5 g | 5 g | 5 g |
| NaCl (Carlo Erba) | 5 g | 5 g | 5 g | 5 g |
| European Bacteriological Agar (Conda) |  | 15 g | 15 g | 15 g |
| Ampicillin (Sigma) (pure) |  |  | 100 mg |  |
| Chloramphenicol (Sigma) (100 mg/ml in ethanol) |  |  | 10 mg |  |
| Cycloheximide (Sigma) (100 mg/ml in DMSO) |  |  |  | 1 mg |

## Supplementary Material 3. Bacterial strains isolated from Oregon-R *Drosophila melanogaster* and used in the experiment

**Figure S3:**
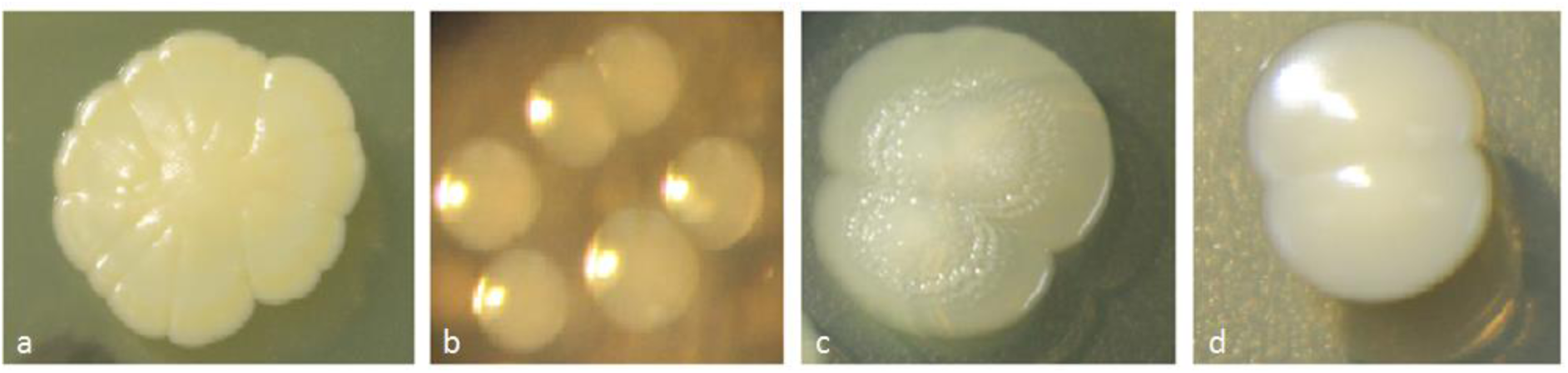
Bacterial strains isolated from Oregon-R *Drosophila melanogaster* and used in the experiment. (a) *Staphylococcus* sp.; (b) *Enterococcus* sp.; (c) Enterobacteriaceae.; (d) Actinobacteria.

## Supplementary Material 4. Experimental design for the grape berry environment

**Figure S4:**
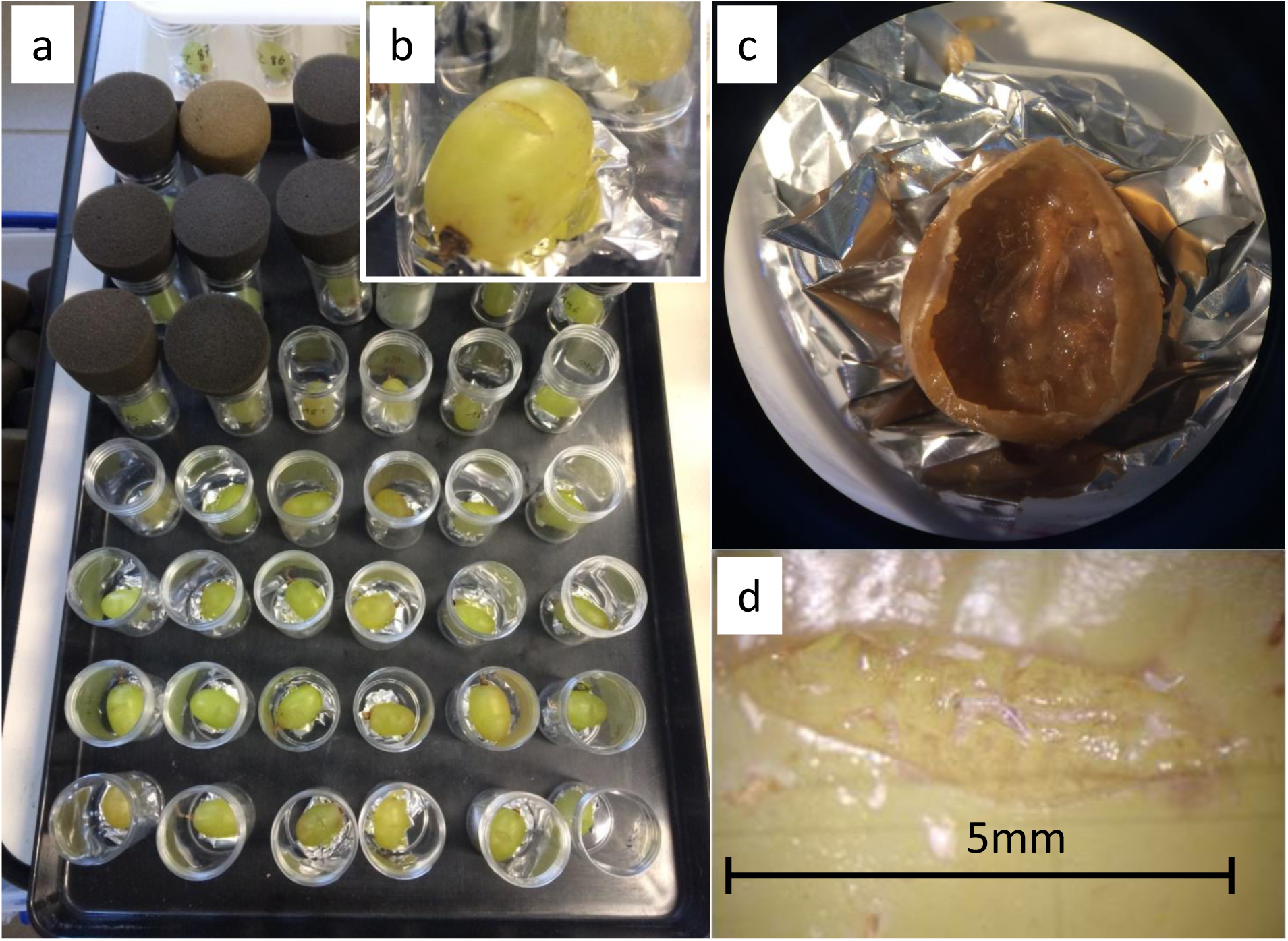
Experimental design for the grape berry environment. (a) Experimental block for grape berry treatments, (b) Experimental unit with grape berry, (c) Decaying grape berry with live yeast, bacteria and larvae, (d) Egg cases visible near berry incision and active larvae in fruit flesh.

## Supplementary Material 5. Bacterial physiological profiles

### Text S5

Eco Microplates (Biolog) were used to have an overview of the metabolic ‘fingerprint’ of the Enterobacteriaceae, the Actinobacteria isolate and the Actinobacteria variant. A fixed number of fresh bacteria cells suspended in sterile PBS were inoculated in well with one of 31 different carbon sources. Each combination Bacterial isolate*Carbon source was replicated three times. The plates were incubated at 25 °C and the absorbance at 595 nm was measured with a Multiskan GO spectrometer (Thermo Scientific) after 48 h and 120 h. A tetrazolium dye included with each carbon source entrained the production of red color when bacterial respiration occurred, i.e. when the carbon source was used. Variations of red color among carbon sources allowed establishing a physiological profile of each bacterial isolate.

**Figure S5.1.**
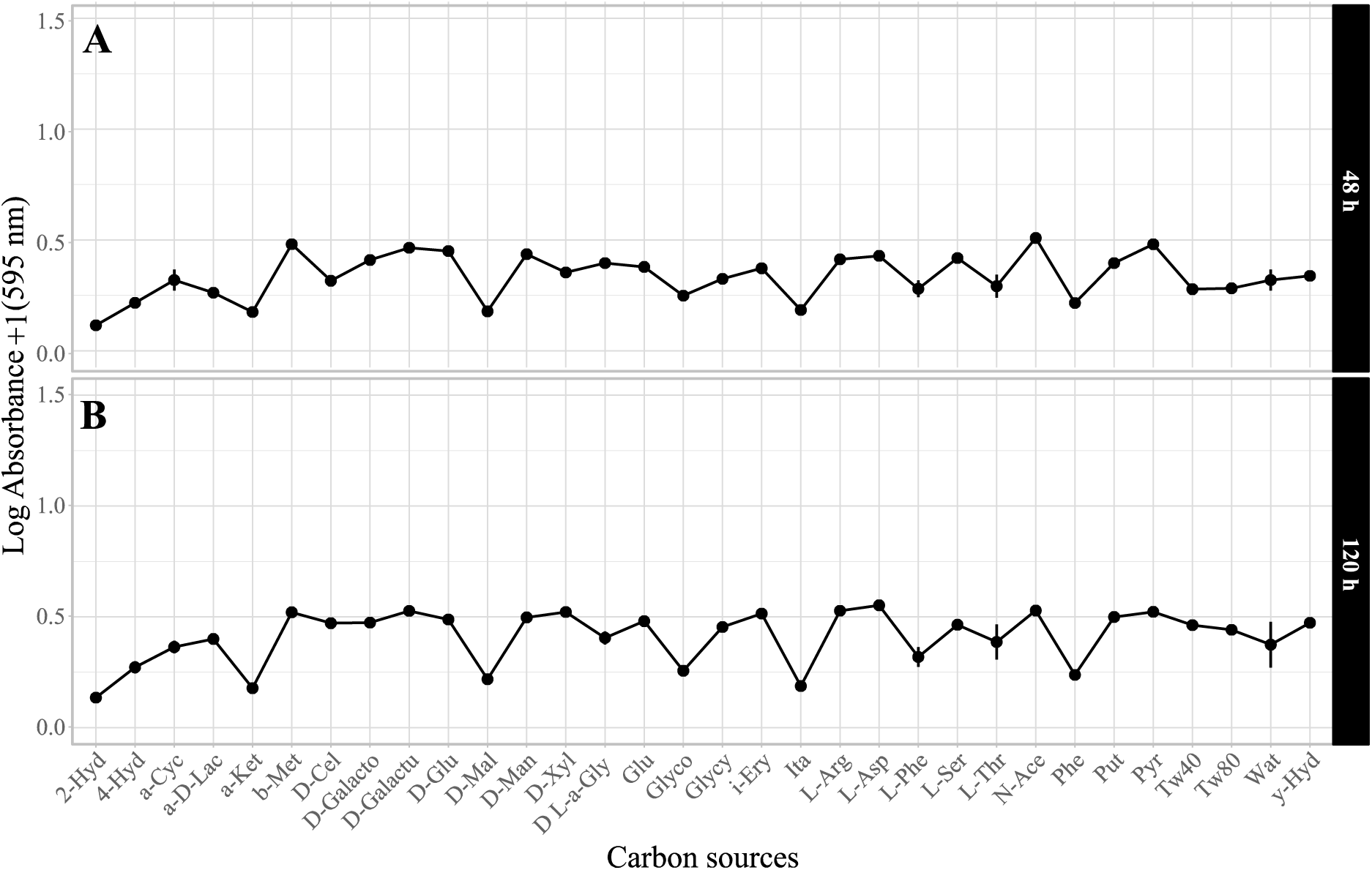
Physiological profile of the Enterobacteriaceae isolate after 48 h- and 120 h-long exposure to different carbon sources. Symbols indicate means; error bars indicate standard errors around the mean. X-axis labels correspond to abbreviations of tested carbon sources, with 2-Hyd for 2-Hydroxy Benzoic Acid; 4-Hyd for 4-Hydroxy Benzoic Acid; a-Cyc for α-Cyclodextrin; a-D-Lac for α-D-Lactose; a-Ket for α-Ketobutyric Acid; b-Met for β-Methyl-D-Glucoside; D-Cel for D-Cellobiose; D-Galacto for D-Galactonic Acid γ-Lactone; D-Galactu for D-Galacturonic Acid; D-Glu for D-Glucosaminic Acid; D-Mal for D-Malic Acid; D-Man for D-Mannitol; D-Xyl for D-Xylose; D L-a-Gly for D,L-α-Glycerol Phosphate; Glu for Glucose-1-Phosphate; Glyco for Glycogen; Glycy for Glycyl-L-Glutamic Acid; i-Ery for i-Erythritol; Ita for Itaconic Acid; L-Arg for L-Arginine; L-Asp for L-Asparagine; L-Phe for L-Phenylalanine; L-Ser for L-Serine; L-Thr for L-Threonine; N-Ace for N-Acetyl-D-Glucosamine; Phe for Phenylethylamine; Put for Putrescine; Pyr for Pyruvic Acid Methyl Ester; Tw40 for Tween 40, Tw80 for Tween 80, Wat for Water and y-Hyd for γ-Hydroxybutyric Acid.

**Figure S5.2.**
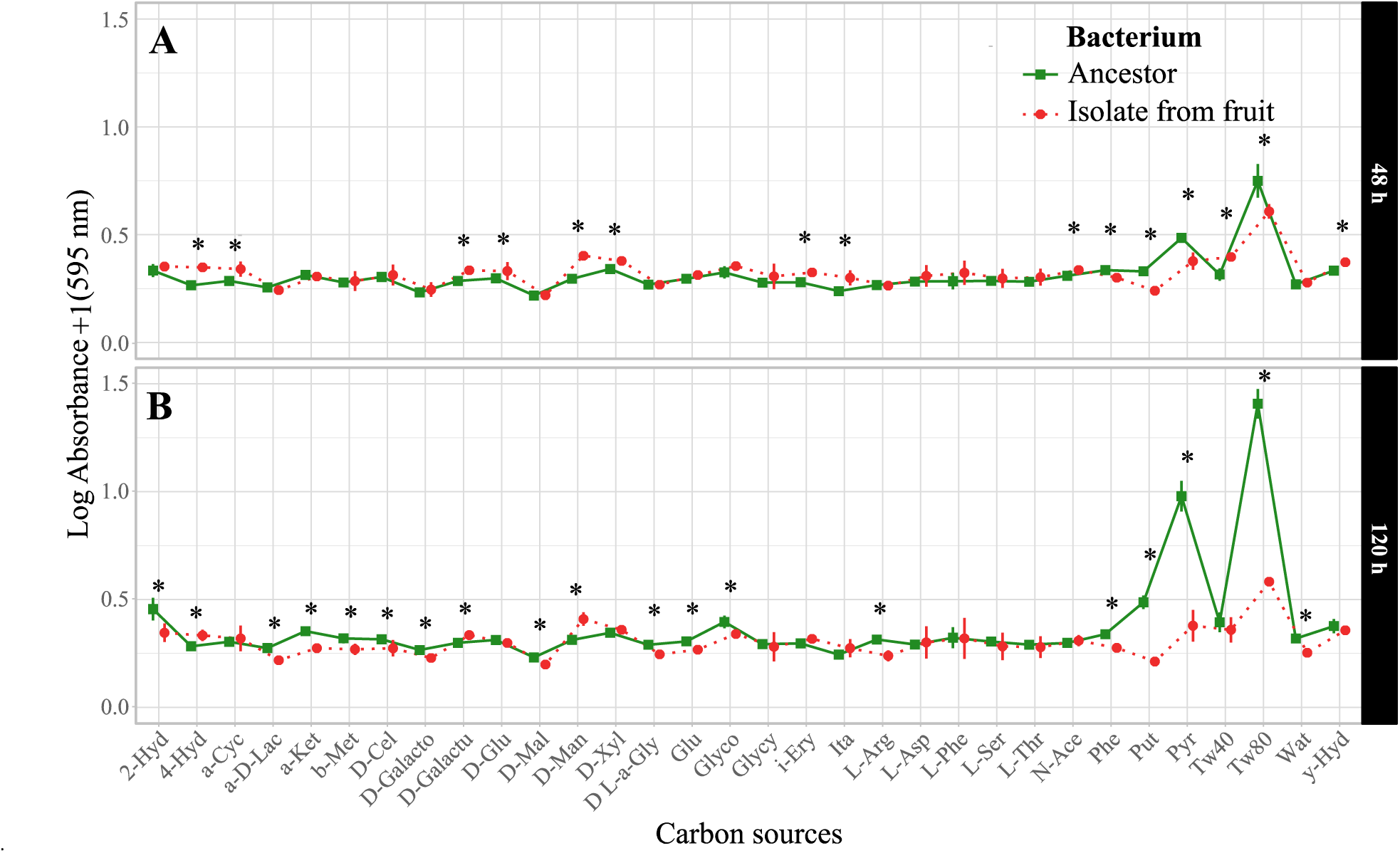
Physiological profiles of the Actinobacteria ancestor and the isolate from grape berry (vial n°419) after 48 h- and 120 h-long exposure to different carbon sources. Symbols indicate means; error bars indicate standard errors around the mean. (*) symbols indicate significant difference between the Actinobacteria ancestor and the isolate from fruit exposed the same duration to a same carbon source. X-axis labels correspond to abbreviations of tested carbon sources, with 2-Hyd for 2-Hydroxy Benzoic Acid; 4-Hyd for 4-Hydroxy Benzoic Acid; a-Cyc for α-Cyclodextrin; a-D-Lac for α-D-Lactose; a-Ket for α-Ketobutyric Acid; b-Met for β-Methyl-D-Glucoside; D-Cel for D-Cellobiose; D-Galacto for D-Galactonic Acid γ-Lactone; D-Galactu for D-Galacturonic Acid; D-Glu for D-Glucosaminic Acid; D-Mal for D-Malic Acid; D-Man for D-Mannitol; D-Xyl for D-Xylose; D L-a-Gly for D,L-α-Glycerol Phosphate; Glu for Glucose-1-Phosphate; Glyco for Glycogen; Glycy for Glycyl-L-Glutamic Acid; i-Ery for i-Erythritol; Ita for Itaconic Acid; L-Arg for L-Arginine; L-Asp for L-Asparagine; L-Phe for L-Phenylalanine; L-Ser for L-Serine; L-Thr for L-Threonine; N-Ace for N-Acetyl-D-Glucosamine; Phe for Phenylethylamine; Put for Putrescine; Pyr for Pyruvic Acid Methyl Ester; Tw40 for Tween 40, Tw80 for Tween 80, Wat for Water and y-Hyd for γ-Hydroxybutyric Acid.

## Supplementary Material 6. Survival on fruit without larvae of the Actinobacteria ancestor and the isolate retrieved from fruit at the end of the experiment

### Text S6

Two concentrations (10,000 and 1,000,000 live cells) of the ancestral Actinobacteria or the Actinobacteria retrieved from the grape berry (vial n°419) suspended in sterile PBS (Phosphate-Buffered Saline) were inoculated on grape slices. The slices of surface-sterilized berries (Behar et al. 2008) were contained in petri dishes with plain agar and incubated at 24 °C. Eight grape slices were sampled and homogenized per treatment after 24 h or 72 h. Numbers of CFUs (Colony Forming Units) were measured on LB agar plates after serial dilutions.

**Figure S6:**
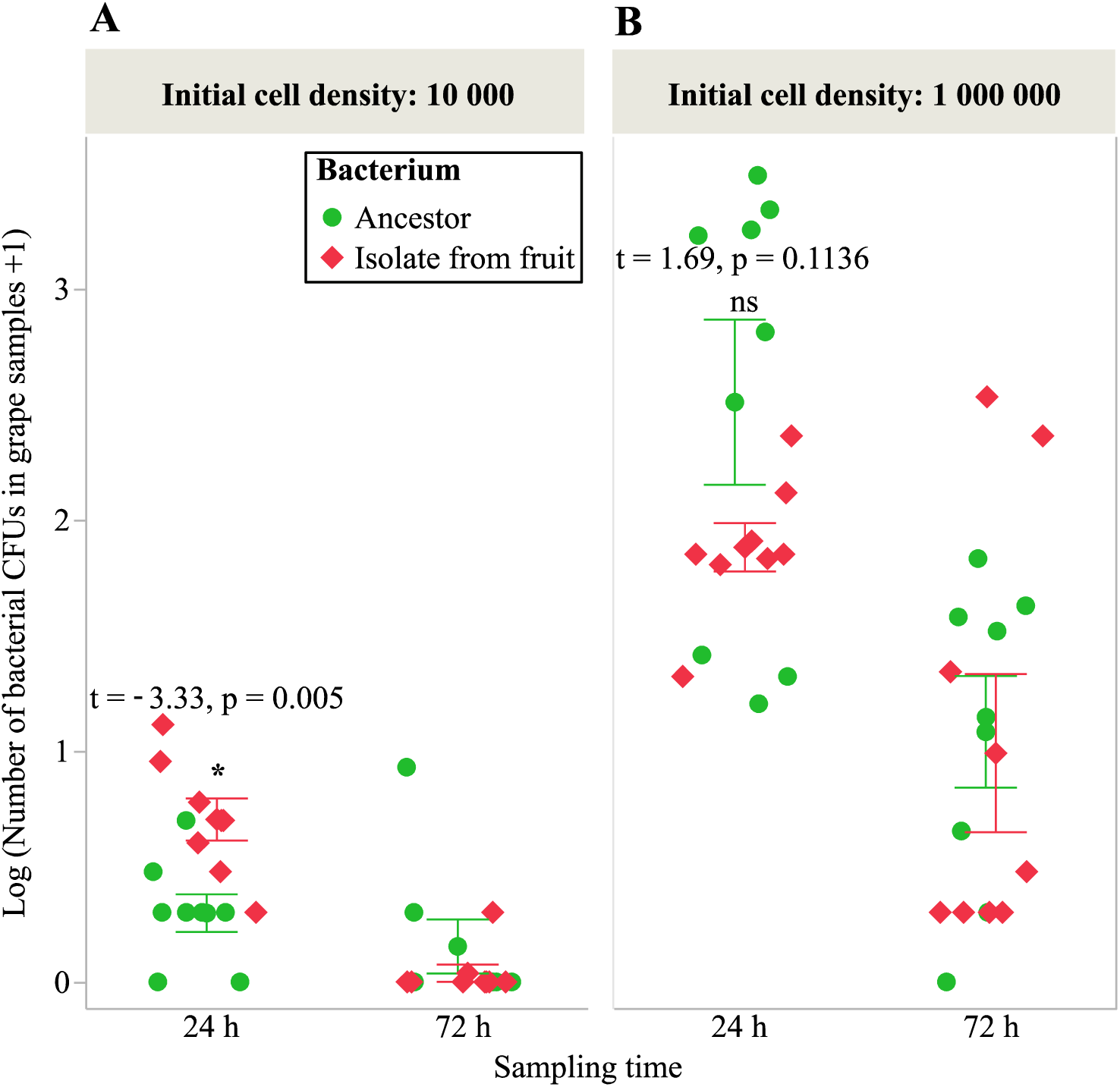
number of CFUs of the Actinobacteria ancestor and the Actinobacteria isolate from grape berry (vial n°419) in samples of grape slices after 24 h- and 72 h-long incubation for two initial cell concentrations. Symbols indicate individual observations; error bars indicate standard errors around the mean. (*) symbol indicates marginally significant difference between the Actinobacteria ancestor and the isolate from fruit.

## Supplementary Material 7. Laboratory medium inoculated with the Enterobacteriaceae

**Figure S7:**
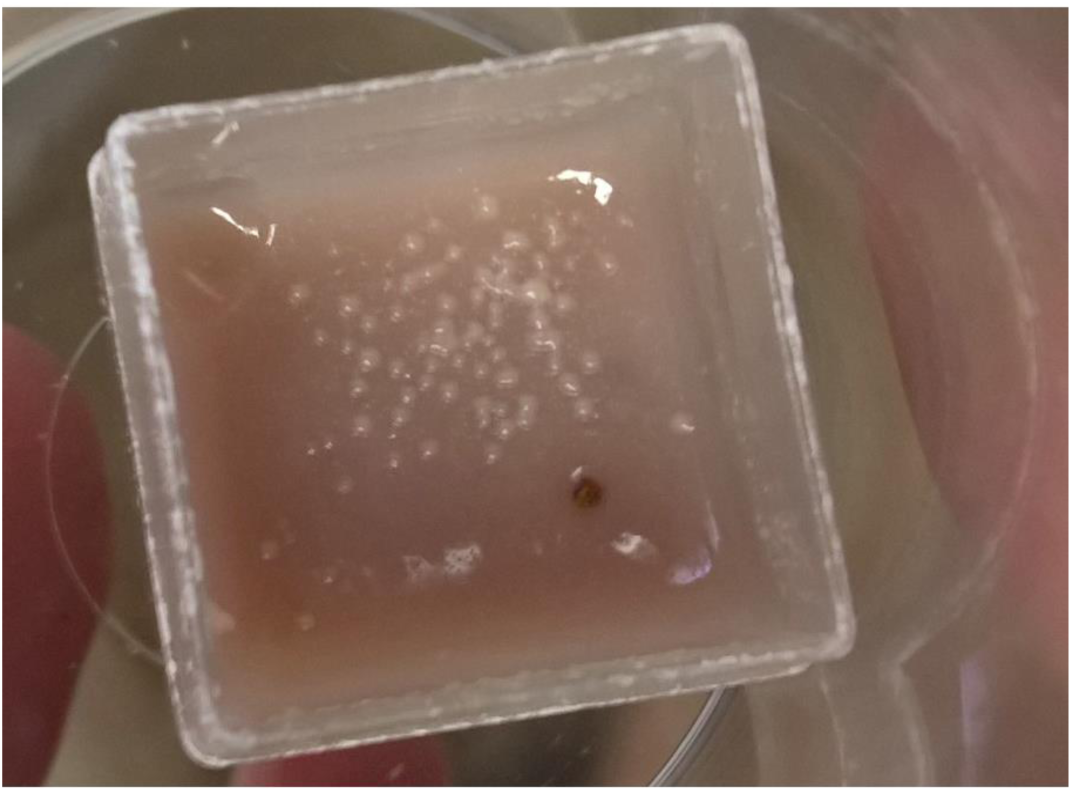
bacterial growth at the surface of laboratory medium five days after Enterobacteriaceae inoculation.

